# c-Maf-dependent regulatory T cells mediate immunological tolerance to intestinal microbiota

**DOI:** 10.1101/129601

**Authors:** Mo Xu, Maria Pokrovskii, Yi Ding, Ren Yi, Christy Au, Carolina Galan, Richard Bonneau, Dan R. Littman

## Abstract

Both microbial and host genetic factors contribute to the pathogenesis of autoimmune disease^1-4^. Accumulating evidence suggests that microbial species that potentiate chronic inflammation, as in inflammatory bowel disease (IBD), often also colonize healthy individuals. These microbes, including the *Helicobacter* species, have the propensity to induce autoreactive T cells and are collectively referred to as pathobionts^4-8^. However, an understanding of how such T cells are constrained in healthy individuals is lacking. Here we report that host tolerance to a potentially pathogenic bacterium, *Helicobacter hepaticus* (*H. hepaticus*), is mediated by induction of RORγt^+^Foxp3^+^ regulatory T cells (iT_reg_) that selectively restrain pro-inflammatory T_H_17 cells and whose function is dependent on the transcription factor c-Maf. Whereas *H. hepaticus* colonization of wild-type mice promoted differentiation of RORγt-expressing microbe-specific iT_reg_ in the large intestine, in disease-susceptible IL-10-deficient animals there was instead expansion of colitogenic T_H_17 cells. Inactivation of c-Maf in the T_reg_ compartment likewise impaired differentiation of bacteria-specific iT_reg_, resulting in accumulation of *H. hepaticus*-specific inflammatory T_H_17 cells and spontaneous colitis. In contrast, RORγt inactivation in T_reg_ only had a minor effect on bacterial-specific T_reg_-T_H_17 balance, and did not result in inflammation. Our results suggest that pathobiont-dependent IBD is a consequence of microbiota-reactive T cells that have escaped this c-Maf-dependent mechanism of iT_reg_-T_H_17 homeostasis.

## Main Text

We chose *Helicobacter hepaticus* (*H*. *hepaticus*) as a model to investigate host-pathobiont interplay. In mice with impaired anti-inflammatory IL-10 signaling, *H*. *hepaticus* induces inflammation of the large intestine (LI), marked by production of interferon gamma (IFNγ) and IL-17^7,8^, with T_H_17 cells accounting for up to 50% of total CD4^+^ T cells (Extended Data Fig. 1a). Such an increase in T_H_17 cells was not observed in the LI of *H*. *hepaticus*-colonized specific pathogen-free (SPF) mice in the absence of IL-10 blockade (Extended Data Fig. 1a). Therefore we wished to determine why *H. hepaticus*-induced T cells do not cause disease in wild-type (WT) animals at steady state. To tackle this question, we initially identified the T cell receptor (TCR) sequences and cognate epitopes of *H. hepaticus*-induced T_H_17 cells present in inflammatory conditions, and subsequently traced the fate of these cells at steady state.

**Figure 1.**
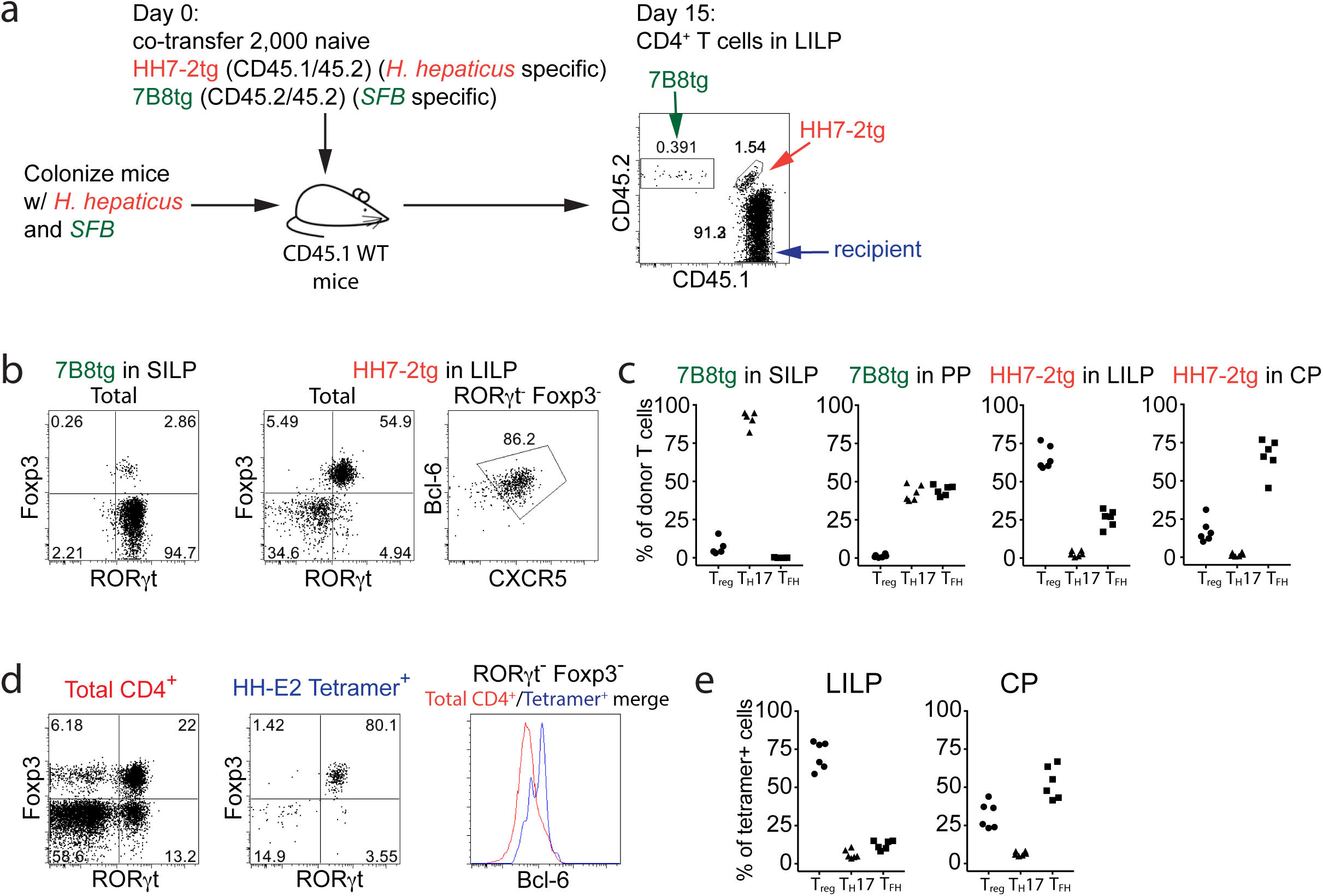
*H. hepaticus* induces RORγt^+^ T_reg_ and T_FH_ responses under steady state conditions. **a**, Experimental scheme for co-transfer and analysis of HH7-2tg and 7B8tg cells in wild type (WT) mice colonized with both *H. hepaticus* and SFB. **b,** Representative flow cytometry plots of RORγt, Foxp3, Bcl-6 and CXCR5 expression in donor-derived T cells in different tissues (n=15). **c,** Frequencies of T_reg_ (Foxp3^+^), T_H_17 (Foxp3^-^RORγt^+^) and T_FH_ (Bcl-6^+^CXCR5^+^) among 7B8tg and HH7-2tg donor cells in indicated tissues. Data are from one of 3 experiments, with total of 15 mice in the 3 experiments. **d,** Representative flow cytometry plots of RORγt, Foxp3 and Bcl-6 expression in total CD4^+^ (red) and HH-E2 tetramer^+^ (blue) T cells from the large intestine of WT mice (n=6) colonized with *H. hepaticus* for 3-4 weeks. **e,** Frequencies of T_reg_ (Foxp3^+^), T_H_17 (Foxp3^-^RORγt^+^) and T_FH_ (Bcl-6^+^) among HH-E2 tetramer^+^ T cells in the indicated tissues of WT mice (n=6) colonized with *H. hepaticus* for 3-4 weeks. Data are a summary of two independent experiments. SILP: small intestinal lamina propria; LILP: large intestinal lamina propria; PP: Peyer's patches and CP: cecal patch.

We sorted GFP^+^ intestinal T_H_17 cells from *H. hepaticus-*colonized *Il23r*^*GFP*^ reporter mice treated with IL-10RA blocking antibody, and performed single cell cloning of T cell receptor (TCR) cDNAs from 384 cells (Extended Data Fig. 1b). We focused on twelve TCR heterodimers that were retrieved from at least two cells. By using a hybridoma carrying NFAT-GFP^+^, which serves as a reporter of TCR signaling, we found that the majority (nine out of twelve) of these TCRs were *H. hepaticus-*specific. We subsequently used a whole-genome shotgun cloning and expression screen^9,10^ to identify a *H. hepaticus-*unique outer membrane protein, HH_1713, as an immunodominant antigen, and further pinpointed two epitopes in HH_1713. The E1 peptide epitope stimulated *H. hepaticus*-specific TCR HH5-1, whereas E2 stimulated TCR HH5-5, HH6-1 and HH7-2 (Extended Data Fig. 1c). We next developed two complementary approaches to track *H. hepaticus*-specific T cells *in vivo*, including HH7-2 and HH5-1 TCR transgenic mice (HH7-2tg and HH5-1tg) ^11,12^ and a MHCII-tetramer loaded with E2 peptide (HH-E2 tetramer) ^13,14^. We validated the specificity of these tracking tools *in vitro* and *in vivo* (Extended Data Fig. 1d-g)

To track what happens to *H. hepaticus*-specific T cells in healthy animals, we simultaneously transferred naïve HH7-2tg and 7B8tg (segmented filamentous bacteria (SFB)-specific TCRtg control)^10^ T cells into WT mice, stably colonized with both *H. hepaticus* and SFB. Donor- and recipient-derived T cells were distinguished using CD45.1 and CD45.2 congenic markers (Fig. 1a). Two weeks post-adoptive transfer, we detected both HH7-2tg and 7B8tg donor T cells in the spleen (SP), mesenteric lymph nodes (mLNs), small and large (including cecum and colon) intestinal lamina propria (SILP and LILP), Peyer's patches (PPs) and the cecal patch (CP) (Extended Data Fig. 2a, b). HH7-2tg donor cells were enriched in the LILP and CP, whereas 7B8tg donor cells predominated in the SILP and PP (Extended Data Fig. 2a, b). The anatomical distribution of the two donor T cell populations was consistent with the colonization of *H. hepaticus* in the LI and SFB in the SI. We next explored the phenotypes of the donor-derived T cells by staining for lineage-specific transcription factors. As previously reported, the vast majority of 7B8 cells developed into T_H_17 cells in the SILP, as they were predominantly positive for RORγt though negative for Foxp3^10^ (Fig. 1b, c). In contrast, HH7-2tg cells in the LILP of WT recipients were mostly iT_reg_ cells that expressed both RORγt and Foxp3 (accounting for ∼60% of total donor-derived HH7-2tg cells)^15,16^, rather than T_H_17 cells (<10% of total HH7-2tg cells) (Fig. 1b, c). There was also a sub-population of 7B8tg and HH7-2tg T cells that expressed neither RORγt nor Foxp3 and predominantly localized to the PPs and CP. The vast majority of these were T follicular helper (T_FH_) cells, marked by co-expression of Bcl-6 and CXCR5 (Fig. 1b, c, Extended Data Fig. 2c, d). When bred onto the *Rag1*^-/-^ background, HH7-2tg mice did not generate T_reg_ in the thymus (Extended Data Fig. 3a), but naïve T cells from these mice differentiated into iT_reg_ when adoptively transferred into *H. hepaticus*-colonized WT mice (Extended Data Fig. 3b, c). These results rule out that HH7-2tg iT_reg_ cells detected after adoptive transfer were derived from thymic T_reg_ contamination or were influenced by the presence of dual TCRs generated by incomplete allelic exclusion of the alpha-chain. Consistent with these results, adoptively transferred HH5-1tg and HH-E2-tetramer positive cells had similar differentiation profiles as HH7-2tg cells (Fig. 1d, e and Extended Data Fig. 4a, b). These results indicate that the host responds to the pathobiont *H. hepaticus* by generating an immunotolerant iT_reg_ response rather than a pro-inflammatory T_H_17 response.

**Figure 2.**
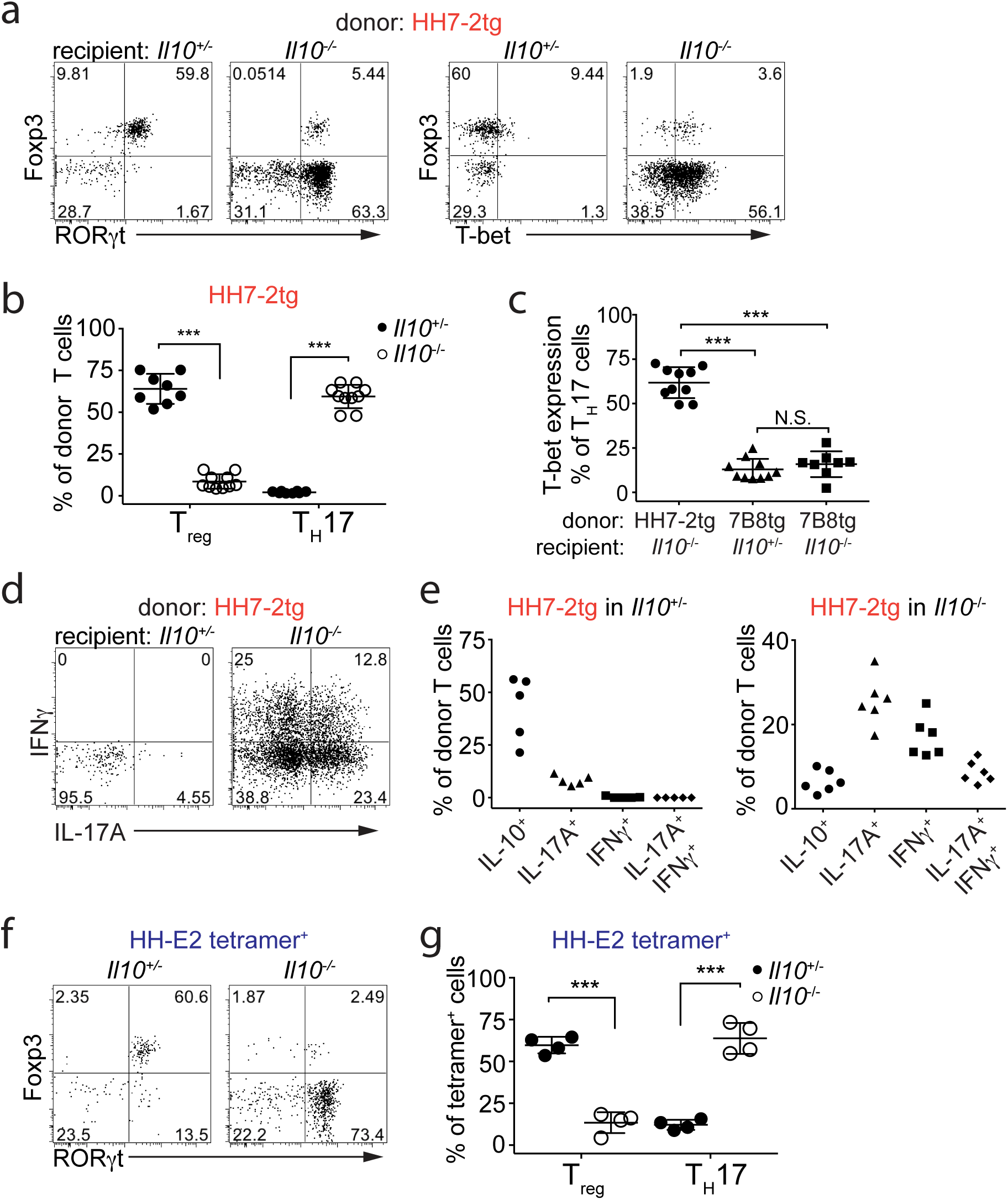
*H. hepaticus* induces inflammatory T_H_17 cells in IL-10 deficiency-dependent colitis. **a,** Representative flow cytometry plots of Foxp3, RORγt and T-bet expression in HH7-2tg donor-derived cells in the LILP of *Il10*^+/-^ (n=8) and *Il10*^-/-^ (n=10) mice. **b,** Frequencies of T_reg_ (Foxp3^+^) and T_H_17 (Foxp3^-^RORγt^+^) cells among LILP HH7-2tg donor-derived cells in *Il10*^+/-^ (n=8) and *Il10*^-/-^ (n=10) mice. Data are from four independent experiments. Error bars: mean ± 1 SD. Statistics were calculated by unpaired *Welch t-*test, *** p<0.001. **c,** Frequencies of T-bet expression among LILP HH7-2tg donor-derived T_H_17 (Foxp3^-^RORγt^+^) cells in *Il10*^-/-^ (n=10) mice, and SILP 7B8tg donor-derived T_H_17 cells in *Il10*^*+/-*^ (n=10) and *Il10*^*-/-*^ (n=8) mice. Data are a summary of four independent experiments. Error bars: mean ± 1 SD. Statistics were calculated by unpaired *Welch t-*test, N.S. (P≥0.05, not significant), *** p<0.001. **d,** Representative flow cytometry plots of IL-17A and IFNγ expression among LILP HH7-2tg donor-derived cells in *Il10*^+/-^ (n=5) and *Il10*^-/-^ (n=6) mice after in vitro re-stimulation. **e,** Frequencies of IL-10, IL-17A and IFNγ expression among LILP HH7-2tg donor-derived cells in *Il10*^+/-^ (n=5) and *Il10*^-/-^ (n=6) mice after re-stimulation. Data are a summary of two independent experiments. **f,** Representative flow cytometry plots of RORγt and Foxp3 expression in HH-E2 tetramer^+^ T cells in the LILP of *Il10*^+/-^ (n=4) and *Il10*^-/-^ (n=4) mice colonized with *H. hepaticus* for 3-4 weeks. **g.** Frequencies of T_reg_ (Foxp3^+^) and T_H_17 (Foxp3^-^RORγt^+^) cells among HH-E2 tetramer^+^ T cells in the LILP of *Il10*^+/-^ (n=4) and *Il10*^-/-^ (n=4) mice colonized with *H. hepaticus* for 3-4 weeks. Data are a summary of two independent experiments. Error bars: mean ± 1 SD. Statistics were calculated by unpaired *Welch t-*test, *** p<0.001.

To examine if the iT_reg_-biased differentiation of *H. hepaticus*-specific T cells is perturbed in the context of intestinal inflammation, we co-transferred naïve HH7-2tg and control 7B8tg T cells into *H. hepaticus*-and SFB-colonized *Il10*^*-/-*^ recipients. Strikingly, unlike in *Il10*^*+/-*^ healthy mice, only a small proportion of transferred HH7-2tg T cells expressed Foxp3 in the LILP of *Il10*^*-/-*^ mice. Instead, most of them differentiated into T_H_17 cells with T_H_1-like features, characterized by expression of both RORγt and T-bet, which was not observed with SFB-specific T_H_17 cells (Fig. 2a-c and Extended Fig. 5a,b). Upon re-stimulation, HH7-2tg cells in control *Il10*^*+/-*^ mice produced IL-10 and very little IL-17A, consistent with the role of Foxp3 in suppressing pro-inflammatory T_H_17 cytokines^17^ (Extended Fig. 5c). By contrast, HH7-2tg cells from *Il10*^*-/-*^ mice produced high levels of both IL-17A and IFNγ, a typical signature of T_H_17-T_H_1 trans-differentiation, which was reported to correlate with inflammation in previous studies^18^ (Fig. 2d, e and Extended Fig. 5c, d). These data were recapitulated with HH-E2 tetramer staining and adoptive transfer of HH5-1tg T cells (Fig. 2f, g and Extended Data Fig. 4c). By comparison, the pro-inflammatory environment in *Il10*^*-/-*^ mice did not result in deviation of SFB-specific T_H_17 cells to the inflammatory T_H_17-T_H_1 phenotype, as these cells neither acquired a higher level of T-bet nor expressed IFNγ following re-stimulation (Extended Data Fig. 5). Therefore, disruption of IL-10-mediated immune tolerance redirected *H. hepaticus*-specific T cells to an inflammatory T_H_17 program, but did not change the fate of SFB-specific T cells.

Finding that *H. hepaticus*-specific T cells are restrained as RORγt^+^ T_reg_ cells in healthy hosts suggested that these cells could be critical for immune tolerance to pathobionts. Yet, the molecular mechanism that maintains this antigen-specific iT_reg_-T_H_17 axis balanced toward T_reg_ remains an open question. The transcription factor c-Maf attracted our attention, because recent studies have found it to be highly enriched in RORγt^+^ iT_reg_^15,19^, and it is known to directly promote RORγt expression while maintaining an anti-inflammatory program, e.g. directing IL-10 expression in other T helper subsets^20,21^. Flow cytometric analysis confirmed c-Maf to be most highly expressed in RORγt^+^ Foxp3^+^ cells compared to other CD4^+^ T cell subsets, including RORγt^-^ Foxp3^+^ cells, in the LI (Fig. 3a). We subsequently deleted *Maf* with *Foxp3*^*cre*^ to test its function in T_reg_. In *Maf*^fl/fl^; *Foxp3*^*cre*^ (*Maf*^Δ*Treg*^) mice there was a marked decrease in the fraction of RORγt^+^ but not RORγt^-^ T_reg_ among CD4^+^ T cells in the LI (Fig. 3b, Extended Data Fig. 6a, c). The remaining small proportion of RORγt^+^ T_reg_ in *Maf*^Δ*Treg*^ mice had residual c-Maf expression, suggesting incomplete protein depletion at the time of analysis (Extended Data Fig. 6b). In addition, *Maf*^Δ*Treg*^ mice exhibited a marked increase in proportions and total numbers of T_H_17 cells, whereas this phenotype was less striking in *Rorc*^*fl/fl*^;*Foxp3*^*cre*^ (*Rorc*^Δ*Treg*^) mice (Fig. 3b). The altered frequency of RORγt^+^ T_reg_ and T_H_17 subsets led us to test if the fate of *H. hepaticus*-specific T cells would be affected in the *Maf*^Δ*Treg*^ and *Rorc*^Δ*Treg*^ mice. Strikingly, HH-E2-tetramer^+^ cells were predominantly T_H_17 in *Maf*^Δ*Treg*^ animals, as compared to being mostly RORγt^+^ T_reg_ in the control mice (Fig. 3c, Extended Data Fig. 6d). In contrast, although *Rorc*^Δ*Treg*^ mice also had increased *H. hepaticus*-specific T_H_17 cells, the majority of tetramer^+^ cells were T_reg_ (Fig. 3c, Extended Data Fig. 6d). Collectively, these results suggest that microbe-specific RORγt^+^ iT_reg_ cells are required for the suppression of inflammatory T_H_17 cell accumulation. Moreover, while RORγt expression contributes to gut iT_reg_ function, c-Maf plays a more substantial role in the differentiation and/or function of these cells.

**Figure 3.**
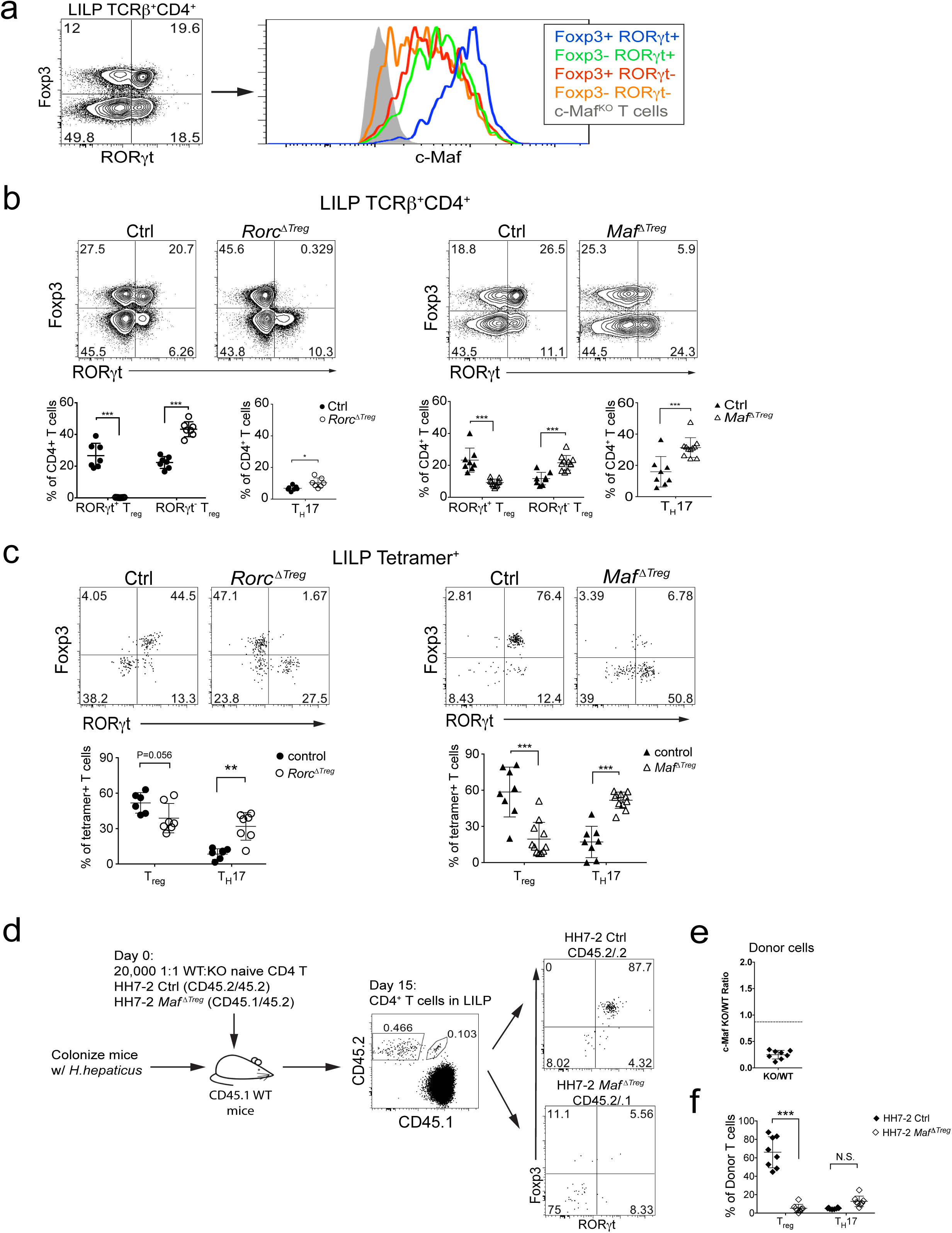
c-Maf is required for the differentiation of induced T_reg_ cells in the gut. **a,** Expression of c-Maf in the indicated CD4^+^ T cell subsets in the LILP. **b.** Transcription factor staining in CD4^+^ T cells from the LILP of mice with T_reg_ cell-specific inactivation of RORγt and c-Maf. Top panels, representative flow cytometry plots of RORγt and Foxp3 expression. Bottom panels, frequencies of RORγt^+^ and RORγt^-^ T_reg_ (Foxp3^+^) cells and T_H_17 (Foxp3^-^RORγt^+^) cells among total CD4^+^ T cells. Data are a summary of 3 independent experiments for *Rorc*^Δ*Treg*^ (n=7) and littermate controls (n=7) and 4 independent experiments for *Maf*^Δ*Treg*^ (n=10) and littermate controls (n=8). Error bars: mean ± 1 SD. Statistics were calculated by unpaired *Welch t-*test, * p<0.05, *** p<0.001. **c,** Transcription factor staining in HH-E2 tetramer positive cells in the LILP. Top panels, representative flow cytometry plots of RORγt and Foxp3 expression. Lower panels, frequencies of T_reg_ (Foxp3^+^) and T_H_17 (Foxp3^-^RORγt^+^) among tetramer positive cells. Data are a summary of 3 independent experiments for *Rorc*^Δ*Treg*^ (n=7) and littermate controls (n=6) and 4 independent experiments for *Maf*^Δ*Treg*^ (n=10) and littermate controls (n=8). Error bars: mean ± 1 SD. Statistics were calculated by unpaired *Welch t-*test, ** p<0.01, *** p<0.001. **d,** Co-transfer of *Maf*^Δ*Treg*^ and control HH7-2tg T cells into WT *H. hepaticus*-colonized mice. Left, schematic of experimental design. Center, representative flow cytometry plots of donor and recipient cell composition in LILP of recipient mice, indicated by CD45.1 and CD45.2. Right, RORγt and Foxp3 expression in *Maf*^Δ*Treg*^ and control HH7-2tg donor-derived cells (n=8). **e,** Ratios of control and *Maf*^Δ*Treg*^ HH7-2tg donor-derived cells (n=8). Dashed line represents ratio of co-transferred cells prior to transfer. Error bars: mean ± 1 SD. **f,** frequencies of T_reg_ (Foxp3^+^) and T_H_17 (Foxp3^-^RORγt^+^) cells among donor-derived cells (n=8). Error bars: mean ± 1 SD. Statistics were calculated by unpaired *Welch t-*test, N.S. (P≥0.05, not significant), *** p<0.001.

We therefore wondered if loss of c-Maf resulted in trans-differentiation of microbe-specific iT_reg_ to T_H_17 cells in a cell intrinsic manner, or if the accumulation of T_H_17 was a consequence of absence of RORγt^+^ iT_reg_ in the tissue. To address this, we co-transferred equal numbers of congenic isotype-labeled naïve *Maf*^*+/+*^*;Foxp3*^*cre*^ and *Maf*^*fl/fl*^*;Foxp3*^*cre*^ HH7-2tg cells into *H. hepaticus*-colonized WT animals. Two weeks after adoptive transfer, the *Maf*^*fl/fl*^*;Foxp3*^*cre*^ HH7-2tg cells were markedly underrepresented compared to control cells in the LI, and were unable to form iT_reg_ (Fig. 3d-f and Extended data Fig. 6e). Importantly, the mutant donor-derived cells did not give rise to a high frequency of T_H_17 cells. Moreover, the vast majority of accumulated T_H_17 cells in *Maf*^Δ*Treg*^ animals expressed c-Maf, indicating that the bulk of these cells did not arise from T_reg_ in which c-Maf was deleted (Extended data Fig. 6f). Thus c-Maf is a critical cell-intrinsic factor for the development of microbe-specific iT_reg_, but suppression of T_H_17 expansion is mediated by these iT_reg_ cells *in trans*.

RORγt expression in iT_reg_ cells has been implicated in the maintenance of gut immune homeostasis under different challenges^15,16^. However, spontaneous gut inflammation in *Rorc*^Δ*Treg*^ animals has not been described. To determine the relative contribution of RORγt or c-Maf in iT_reg_ cells towards immune tolerance, we examined *Rorc*^Δ*Treg*^ and *Maf*^Δ*Treg*^ mice for signs and symptoms of inflammation. We noticed that *Maf*^Δ*Treg*^, but not *Rorc*^Δ*Treg*^ or control littermates, were prone to rectal prolapse (Fig. 4a). Moreover, five to six weeks after colonization with *H. hepaticus*, *Maf*^Δ*Treg*^ mice had enlarged LI-draining mesenteric lymph nodes (mLNs) and increased cellularity of mLNs and LI (Fig. 4b, c). Histopathological analysis of the large intestine of these animals revealed mixed acute and chronic inflammation including multifocal or diffuse mononuclear inflammatory infiltrates in the lamina propria, crypt abscesses or cryptitis and architectural glandular disarray with reactive epithelial changes (nuclear enlargement, mitotic activity, reduced mucin) (Fig. 4d). Notably, none of the above changes was observed in *Rorc*^Δ*Treg*^ mice (Fig. 4b-d, Extended data Fig. 6c, g). Thus, c-Maf but not RORγt expression in iT_reg_ cells is critical for suppression of spontaneous inflammation.

**Figure 4:**
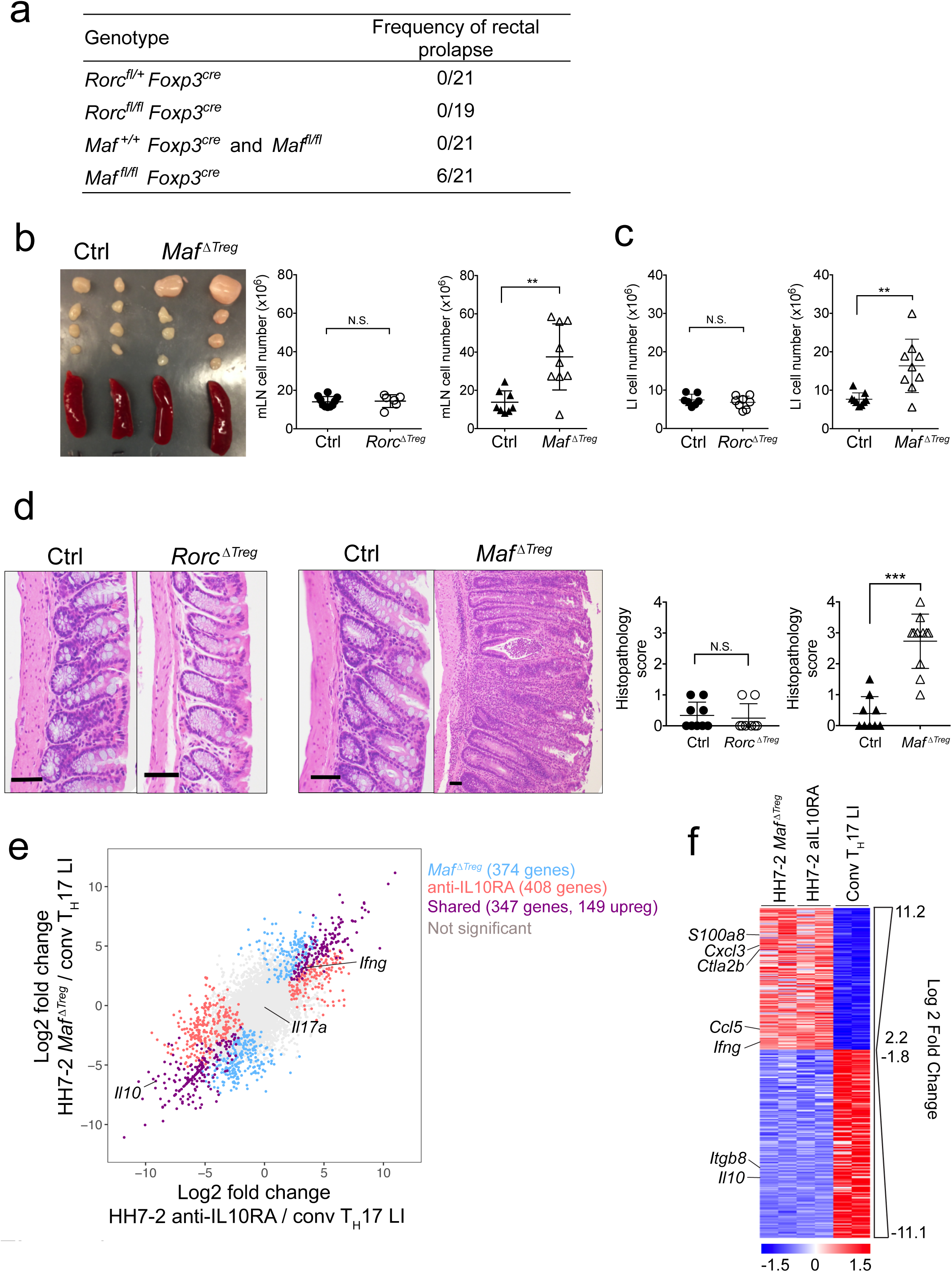
Deletion of c-Maf in T_reg_ cells leads to spontaneous colitis. **a,** Table representing frequency of rectal prolapse by genotype. **b,** Spleens and mesenteric lymph nodes from *Maf*^Δ*Treg*^ and littermate controls (left). Total cell numbers in mLNs (right). Data are a summary of 3 independent experiments for *Rorc*^Δ*Treg*^ (n=6) and littermate controls (n=10), and 4 independent experiments for *Maf*^Δ*Treg*^ (n=9) and littermate controls (n=8). Error bars: mean ± 1 SD. Statistics were calculated by unpaired *Welch t-test*, N.S. (P≥0.05, not significant), ** p<0.01. **c,** Number of leukocytes in the LILP. Data are a summary of 3 independent experiments for *Rorc*^Δ*Treg*^ (n=7) and littermate controls (n=8), and 4 independent experiments for *Maf*^Δ*Treg*^ and littermate controls (n=9). Error bars: mean ± 1 SD. Statistics were calculated by unpaired *Welch t-*test, N.S. (P≥0.05, not significant), ** p<0.01. **d,** Representative histology of large intestine sections from mice with indicated genotypes, colonized with *H. hepaticus* for 4-6 weeks. Scale bar represents 50 µm. Right, colitis scores (0-4 scale) in mice with T_reg_-specific inactivation of transcription factors. *Rorc*^Δ*Treg*^ (n=8) and littermate controls (n=9). *Maf*^Δ*Treg*^ (n=11) and littermate controls (n=9). Error bars: mean ± 1 SD. Statistics were calculated by unpaired *Welch t-*test, N.S. (P≥0.05, not significant), *** P<0.001. **e,** Comparison of transcriptomes of *H. hepaticus*-specific T_H_17 cells from mice treated with IL-10Ra blockade or *Maf*^Δ*Treg*^ and conventional T_H_17 cells (predominantly SFB-specific). Scatter plot depicting log fold change of gene expression. Blue, red and purple dots indicate significant difference (FDR < 0.1). **f,** Heatmap depicting the 347 shared genes differentially expressed (Pval<0.1) between pathogenic HH7-2 and conventional T_H_17 cells. Data for each condition are the mean of 2 biological replicates. Heatmap depicting the 347 shared differentially expressed genes between pathogenic HH7-2 and conventional T_H_17 (purple dots in Fig 4e). Scale bar represents z-scored variance stabilized data (VSD) counts. Data for each condition is the mean of 2 biological replicates.

Because, *Maf*^Δ*Treg*^ and *Il10*^*-/-*^ mice have a similar *H. hepaticus*-driven colitis phenotype, we wished to determine if bacteria-specific T_H_17 cells share an inflammatory regulatory program. We therefore compared the transcriptional profiles of *H. hepaticus*-specific T effector (T_Eff_) cells from *HH7-2tg;Maf*^Δ*Treg*^ mice with spontaneous colitis and *HH7-2tg;Foxp3*^*cre*^ mice with IL-10RA blockade-induced colitis to homeostatic IL-23R-GFP^+^ T cells (which are predominantly SFB-specific T_H_17 cells) (Extended Data Fig. 7a). The gene expression profiles of T_Eff_ from the *Maf*^Δ*Treg*^ and anti-IL10RA-treated HH7-2tg animals were distinct from those of homeostatic T_H_17 cells, as shown by principal component analysis (PCA) (Extended data Fig. 7b, c). Comparison of HH7-2tg T_Eff_ cells to homeostatic T_H_17 cells revealed 1,129 differentially expressed genes, 149 of which were up-regulated in T_Eff_ from both *Maf*^Δ*Treg*^ and anti-IL10RA-treated mice (Fig. 4e, Extended data Fig. 7e). Further analysis of this shared gene set revealed transcripts associated with pathogenic T_H_17 cells (*Ifng, A100a8, Cxcl3, Ccl5, Ctla2b*) and inflammatory disease pathways (Fig. 4f, Extended data Fig. 7d). These data indicate that *H.hepaticus*-specific T_H_17 cells in *Maf*^Δ*Treg*^ mice are highly similar to pathogenic T_H_17 cells in *Il10*^*-/-*^ mice, but differ markedly from homeostatic T_H_17 cells.

As c-Maf is also expressed, albeit at a lower level, in thymus-derived nT_reg_ cells, we considered whether it also has an important role in the function of these cells. We sorted Neuropilin-1^+^ (NPR1^+^) nT_reg_ from the spleen and peripheral lymph nodes (pLNs) of *Maf*^Δ*Treg*^ mice and their control littermates, and evaluated their functions^22,23^. C-Maf-deficient and –sufficient nT_reg_ showed equal activity in inhibiting T_Eff_ cell proliferation in vitro, as well as in suppressing pathogenesis in a model of T cell transfer colitis *in vivo* (Extended Data Fig. 8). These results argue against the possibility that a defect in nT_reg_ cells contributes to the accumulation of inflammatory T_H_17 cells and spontaneous development of colitis in *Maf*^Δ*Treg*^ mice, further highlighting the critical function of iT_reg_ in the maintenance of tolerance to gut pathobionts.

Our results reveal a mechanism for how a common commensal pathobiont like *H. hepaticus* can co-exist with a healthy host without causing disease. Through induction of tolerogenic c-Maf-dependent iT_reg_, the immune system constrains pro-inflammatory *H. hepaticus*-specific T_H_17 cells. Our results are consistent with and help explain the expansion of colitogenic T_H_17 cells in mice with T_reg_-specific inactivation of Stat3^24^. Like c-Maf, Stat3 is likely required for the differentiation and/or function of microbiota-induced RORγt^+^ iT_reg_ cells. The accumulation of similar colitogenic T_H_17 cells in IL-10 signaling-deficient and *Maf*^Δ*Treg*^ mice is consistent with the role of RORγt^+^ T_reg_ cells as the most robust IL-10 producing T cell subset^25^. This work represents a significant step toward elucidating the mechanisms by which the host immune system contains inflammatory disease induced by pathobionts. Our results also raise the question of why benign SFB-induced T_H_17 responses are not constrained by iT_reg_ cells while they suggest a mechanism whereby commensalism is established by balancing induction of microbe-specific iT_reg_ and inflammatory Th17 cells, with the regulatory cells keeping inflammation at bay.

## METHODS

### Mice

Mice were bred and maintained in the animal facility of the Skirball Institute (New York University School of Medicine) in specific pathogen-free conditions. C57Bl/6 mice were obtained from Jackson Laboratories or Taconic Farm. *Il10*^-/-^ (B6.129P2-*Il10*^*tm1Cgn*^/J) mice were purchased from Jackson Laboratories and bred with WT C57Bl/6 mice, which subsequently generated *Il10*^+/-^ and *Il10*^-/-^ littermates by heterozygous breeding. *CD45.1* (*B6.SJL-Ptprca Pepcb/BoyJ*) mice were purchased from Jackson Laboratories. *Foxp3*^*creYFP*^ mice were previously described and obtained from Jackson Laboratories^26^. *Il23r*^*gfp*^ and *Maf*^*fl/fl*^ strains were previously described^27,28^ and kindly provided by Drs. M. Oukka and C. Birchmeier, respectively. All animal procedures were performed in accordance with protocols approved by the Institutional Animal Care and Usage Committee of New York University School of Medicine.

### Antibodies, intracellular staining and flow cytometry

The following monoclonal antibodies were purchased from eBiosciences, BD Pharmingen or BioLegend: CD3 (145-2C11), CD4 (RM4-5), CD25 (PC61), CD44 (IM7), CD45.1 (A20), CD45.2 (104), CD62L (MEL-14), CXCR5 (L138D7), NPR-1 (3E12), TCRβ (H57-597), TCR Vβ6 (RR4-7), TCR Vβ8.1/8.2 (MR5-2), TCR Vβ14 (14-2), Bcl-6 (K112-91), c-Maf (T54-853), Foxp3 (FJK-16s), GATA3 (TWAJ), Helios (22F6), RORγt (B2D or Q31-378), T-bet (eBio4B10), IL-10 (JES5-16E3), IL-17A (eBio17B7) and IFN-γ (XM61.2). 4',6-diamidino-2-phenylindole (DAPI) or Live/dead fixable blue (ThermoFisher) was used to exclude dead cells.

For transcription factor staining, cells were stained for surface markers, followed by fixation and permeabilization before nuclear factor staining according to the manufacturer's protocol (Foxp3 staining buffer set from eBioscience). For cytokine analysis, cells were incubated for 5 h in RPMI with 10% FBS, phorbol 12-myristate 13-acetate (PMA) (50 ng/ml; Sigma), ionomycin (500 ng/ml; Sigma) and GolgiStop (BD). Cells were stained for surface markers before fixation and permeabilization, and then subjected to intracellular cytokine staining according to the manufacturer's protocol (Cytofix/Cytoperm buffer set from BD Biosciences).

Flow cytometric analysis was performed on an LSR II (BD Biosciences) or an Aria II (BD Biosciences) and analyzed using FlowJo software (Tree Star).

### Isolation of lymphocytes

Intestinal tissues were sequentially treated with PBS containing 1 mM DTT at room temperature for 10 min, and 5 mM EDTA at 37°C for 20 min to remove epithelial cells, and then minced and dissociated in RPMI containing collagenase (1 mg/ml collagenase II; Roche), DNase I (100 µg/ml; Sigma), dispase (0.05 U/ml; Worthington) and 10% FBS with constant stirring at 37°C for 45 min (SI) or 60 min (LI). Leukocytes were collected at the interface of a 40%/80% Percoll gradient (GE Healthcare). The Peyer's patches and cecal patch were treated in a similar fashion except for the first step of removal of epithelial cells. Lymph nodes and spleens were mechanically disrupted.

### Single-cell TCR cloning

*Il23r*^*GFP/+*^ mice were maintained in SFB-free conditions to guarantee low T_H_17 background levels. To induce robust T_H_17 response, the mice were orally infected with *H. hepaticus* and injected intraperitoneally with 1mg anti-IL10RA (clone 1B1.3A, Bioxcell) every week from the day of infection. After two weeks, LI GFP^+^ CD4^+^ T cells were sorted on the BD Aria II and deposited at one cell per well into 96-well PCR plates pre-loaded with 5 µl high-capacity cDNA reverse transcription mix (Thermo Fisher) supplemented with 0.1% Triton X-100 (Sigma-Aldrich). Immediately after sorting, whole plates were incubated at 37 °C for 2 h, and then inactivated at 85 °C for 10 min for cDNA. A nested multiplex PCR approach described previously was used to amplify the CDR3α and CDR3β TCR regions separately from the single cell cDNA^29^. PCR products were cleaned up with ExoSap-IT reagent (USB) and Sanger sequencing was performed by Macrogen. Open reading frame nucleotide sequences of the TCRα and TCRβ families were retrieved from the IMGT database (http://www.imgt.org)^30^.

### Generation of TCR hybridomas

The NFAT-GFP 58α^-^β^-^ hybridoma cell line was kindly provided by Dr. K. Murphy^31^. To reconstitute TCRs, cDNA of TCRα and TCRβ were synthesized as gBlocks fragments by Integrated DNA Technologies (IDT), linked with the self-cleavage sequence of 2A (TCRα-p2A-TCRβ), and shuttled into a modified MigR1 retrovector in which IRES-GFP was replaced with IRES-mCD4 (mouse CD4) as described previously^10^. Then retroviral vectors were transfected into Phoenix E packaging cells using TransIT-293 (Mirus). Hybridoma cells were transduced with viral supernatants in the presence of polybrene (8µg/ml) by spin infection for 90 min at 32 °C. Transduction efficiencies were monitored by checking mCD3 surface expression three days later.

### Assay for hybridoma activation

Splenic dendritic cells were used as antigen presenting cells (APCs). B6 mice were injected intraperitoneally with 5 × 10^6^ FLT3L-expressing B16 melanoma cells to drive APC proliferation as previously described^32^. Splenocytes were prepared 10 days after injection, and positively enriched for CD11c^+^ cells using MACS LS columns (Miltenyi). 2 × 10^4^ hybridoma cells were incubated with 10^5^ APCs and antigens in round bottom 96-well plates for two days. GFP induction in the hybridomas was analyzed by flow cytometry as an indicator of TCR activation.

### Construction and screen of whole-genome shotgun library of *H. hepaticus*

The shotgun library was prepared with a procedure modified from previous studies^9,10^. In brief, genomic DNA was purified from cultured *H. hepaticus* with DNeasy PowerSoil kit (Qiagen). DNA was partially digested with MluCI (NEB), and the fraction between 500 and 2000 bp was ligated into the EcoRI-linearized pGEX-6P-1 expression vector (GE Healthcare). Ligation products were transformed into ElectroMAX DH10B competent Cells (Invitrogen) by electroporation. To estimate the size of the library, we cultured 1% and 0.1% of transformed bacteria on lysogeny broth (LB) agar plates containing 100µg/mL Ampicillin for 12 h and then quantified the number of colonies. The library is estimated to contain 3X10^4^ clones. To ensure the quality of the library, we sequenced the inserts of randomly picked colonies. All the sequences were mapped to the *H. hepaticus* genome, and their sizes were 700 to 1200 bp. We aliquoted the bacteria into 96-well deepwell plates (Axygen) (∼30 clones/well) and grew with AirPort microporous cover (Qiagen) in 37°C. The expression of exogenous proteins was induced by 1mM isopropylthiogalactoside (IPTG, Sigma) for 4 h. Then bacteria were collected in PBS and heat-killed by incubating at 85 °C for 1 h, and stored at -20 °C until use. Two screening rounds were performed to identify the antigen-expressing clones. For the first round, pools of heat-killed bacterial clones were added to a co-culture of splenic APCs and hybridomas. Clones within the positive pools were subsequently screened individually against the hybridoma bait. Finally, the inserts of positive clones were subjected to Sanger sequencing. The sequences were blasted against the genome sequence of *H. hepaticus* (ATCC51449) and aligned to the annotated open reading frames. Full-length open reading frames containing the retrieved fragments were cloned into pGEX-6P-1 to confirm their activity in the T cell stimulation assay.

### Epitope mapping

We cloned overlapping fragments spanning the entire HH_1713 coding region into the pGEX-6P-1 expression vector, and expressed these in *E. coli* BL21 cells. The heat-killed bacteria were used to stimulate relevant hybridomas. This process was repeated until we mapped the epitope to a region containing 30 amino acids. The potential MHCII epitopes were predicted with online software RANKPEP^33^. Overlapping peptides spanning the predicted region were further synthesized (Genescript) and verified by stimulation of the hybridomas.

### Generation of TCRtg mice

TCR sequences of HH5-1 and HH7-2 were cloned into the pTα and pTβ vectors kindly provided by Dr. D. Mathis. TCR transgenic animals were generated by the Rodent Genetic Engineering Core at the New York University School of Medicine. Positive pups were genotyped by testing TCR Vβ8.1/8.2 (HH5-1tg) or Vβ6 (HH7-2tg) expression on T cells from peripheral blood.

### MHCII tetramer production and staining

HH-E2 tetramer was kindly produced by the NIH Tetramer Core Facility. Briefly, QESPRIAAAYTIKGA (HH_1713-E2), an immunodominant epitope validated with the hybridoma stimulation assay, was covalently linked to I-A^b^ via a flexible linker, to produce pMHCII monomers. Soluble monomers were purified, biotinylated, and tetramerized with phycoerythrin- or allophycocyanin-labelled streptavidin. To stain endogenous T cells, mononuclear cells from LILP or CP were first resuspended in MACS buffer with FcR block, 2% mouse serum and 2% rat serum. Then tetramer was added (10 nM) and incubated at room temperature for 60 min, and cells were re-suspended by pipetting every 20 min. Cells were washed with MACS buffer and followed by regular surface marker staining at 4 °C.

### Adoptive transfer of TCRtg cells

Spleens from TCRtg mice were collected and mechanically disassociated. Red blood cells were lysed using ACK lysis buffer (Lonza). For TCRtg mice in WT background, naive Tg T cells were sorted as CD4^+^CD3^+^CD44^lo^CD62L^hi^CD25^-^ Vβ6^+^ (HH7-2tg), Vβ8.1/8.2^+^ (HH5-1tg) or Vβ14^+^ (7B8tg) on the Aria II (BD Biosciences). For HH7-2tg mice bred to the *Foxp3*^*creYFP*^ background, naive Tg T cells were sorted as CD4^+^CD3^+^CD44^lo^CD62L^hi^Foxp3^creYFP-^Vβ6^+^. Cells were resuspended in PBS and transferred into congenic isotype-labeled recipient mice by retro-orbital injection.

### *H. hepaticus* culture and oral infection

*H. hepaticus* was kindly provided by Dr. James Fox (MIT). Frozen stock aliquots of *H. hepaticus* were stored in Brucella broth with 20% glycerol and frozen at -80°C. The bacteria were grown on blood agar plates (TSA with 5% sheep blood, Thermo Fisher). Inoculated plates were placed into a hypoxia chamber (Billups-Rothenberg), and anaerobic gas mixture consisting of 80% nitrogen, 10% hydrogen, and 10% carbon dioxide (Airgas) was added to create a micro-aerobic atmosphere, in which the oxygen concentration was 3∼5%. The micro-aerobic jars containing bacterial plates were left at 37°C for 5 days before animal inoculation. For oral infection, *H. hepaticus* was resuspended in Brucella broth by application of a pre-moistened sterile cotton swab applicator tip to the colony surface. The concentration of bacterial inoculation dose was determined by the use of a spectrophotometric optical density (OD) analysis at 600 nm, and adjusted to OD_600_ readings between 1 and 1.5. 0.2 mL bacterial suspension was administered to each mouse by oral gavage. Mice were inoculated every 5 days for a total of two doses.

### T_reg_ cell *in vitro* suppression assay

Naïve T cells (T_naive_) with the phenotype CD4^+^CD3^+^CD44^lo^CD62L^hi^CD25^-^ were isolated from CD45.1 WT B6 mice by FACS and labeled with carboxyfluorescein diacetate succinimidyl ester (CFSE). nT_reg_ (CD45.2) with the phenotype CD4^+^CD3^+^Foxp3^creYFP+^NRP1^+^ were isolated from *Foxp3*^*creYFP*^ or *Maf*^Δ*Treg*^ mice by FACS. B cells were isolated as APCs by positive enrichment for B220^+^ cells using MACS LS columns (Miltenyi) from CD45.2 WT B6 mice. 2.5 × 10^4^ CFSE-labeled T_naive_ cells were cultured for 72 h with APCs (5 ×10^4^) and anti-CD3 (1 µg/ml) in the presence or absence of various numbers of T_reg_ cells. The cell division index of responder T cells was assessed by dilution of CFSE using FlowJo software (Tree Star).

### Adoptive transfer colitis

CD4^+^CD3^+^CD25^-^CD45RB^hi^ T_Eff_ cells were isolated by FACS from B6 mouse spleens. CD4^+^CD3^+^Foxp3^creYFP+^NPR1^+^ nT_reg_ were isolated from the splenocytes of *H. hepaticus*-colonized *Foxp3*^*creYFP*^ or *Maf*^Δ*Treg*^ mice. T_Eff_ cells (5 ×10^5^) were administered by retro-orbital injection into *H. hepaticus*-colonized Rag1^-/-^ mice alone, or simultaneously with 4 ×10^5^ nT_reg_ as previously described^34^. Co-housed littermate recipients were randomly assigned to different treatment groups such that each cage contained all treatment conditions. Animal weights were measured weekly. After eight weeks, large intestines were collected and fixed with 10% neutral buffered formalin (Fisher). Samples were sectioned and stained with Haemotoxylin and Eosin by the Histopathology Core at the New York University School of Medicine.

### Histology analysis

The H&E slides from each sample were examined in a blinded fashion. Samples of proximal, mid, and distal colon were graded semiquantitatively from 0 to 4 as described previously^35^. Scores from proximal, mid, and distal sites were averaged to obtain inflammation scores for the entire colon.

### Cell isolation for RNA-seq experiment

To induce colitis, both *HH7-2tg;Maf* ^Δ*Treg*^ and *HH7-2tg;Foxp3*^*cre*^ mice were colonized with *H. hepaticus*. *HH7-2tg;Foxp3*^*cre*^ mice were further I.P. injected with 1mg anti-IL10RA (clone 1B1.3A, Bioxcell) antibody weekly from the day of colonization. HH7-2 T effector T_Eff_ cells (CD3^+^CD4^+^TCRVβ6^+^Foxp3-YFP^-^) were sorted from the LILP two weeks after colonization. Homeostatic IL-23R-GFP^+^ T cells (CD3^+^CD4^+^IL-23R-GFP^+^) were sorted from both SILP and LILP of *Il23r*^gfp/+^ mice stably colonized with SFB.

### RNA-seq library preparation

Total RNA was extracted using TRIzol (Invitrogen) followed by DNase I treatment and cleanup with RNeasy MinElute kit (Qiagen). RNAseq libraries were prepared using Nugen Ovation Ultralow Library Systems V2 (cat # 7102 and 0344) and sequenced on the Illumina NextSeq.

### Data processing of RNA-seq experiment

RNA-seq reads were mapped to the *Mus musculus* genome Ensembl annotation release 87 with STAR (v2.5.2b)^36^. Uniquely mapped reads were counted using featureCounts^37^ with parameters: -p -Q 20. DESeq2^38^ was used to identify differentially expressed genes across conditions with experimental design: ∼Condition + Gender. Read counts were normalized and transformed by function VarianceStabilizingTransformation (VST) in DESeq2 with the following parameter: blind=FALSE. Gender differences were considered as batch effect, and were corrected by ComBat^39^. Downstream analysis and data visualization were performed in R^40^.

### Statistical analysis

For animal studies, two-sided *Welch t-test* with Holm-Sidak correction for multiple comparisons was used. Error bars represent +/- 1 standard deviation. Final sample sizes matched predicted sample sizes calculated using power analysis. No samples were excluded from analysis. Variance between KO samples tended to be greater than controls. For RNA-seq analysis, genes were considered differentially expressed when DESeq2^38^ adjusted P values were < 0.1. Enriched disease pathways were determined using Ingenuity Pathway Analysis (www.Ingenuity.com).

## Data Availability

cDNA sequences of *H. hepaticus*-specific TCRs data that support the findings of this study have been deposited in GenBank with the accession codes KY964547-KY964570. RNA-seq data that support the findings of this study can be accessed here: http://voms.simonsfoundation.org:50013/hywulth4D4N0XPuO7vsLY7ea04gk19f/cMaf_fastq

## Acknowledgements

We thank S.Y. Kim and the NYU Genome Engineering Core for generating TCR transgenic mice, A. Heguy and colleagues at the NYU School of Medicine's Genome Technology Center (GTC) for timely preparation of RNA-seq libraries and RNA-sequencing, the NIH Tetramer Core Facility for generating MHC II tetramers, K. Murphy for providing the 58α-β- hybridoma line, P. Dash and P.G. Thomas for suggestions on single cell TCR cloning, and J.A. Hall, J. Muller and J. Lafaille for suggestions on the manuscript. The Experimental Pathology Research Laboratory of NYU Medical Center is supported by National Institutes of Health Shared Instrumentation Grants S10OD010584-01A1 and S10OD018338-01. The GTC is partially supported by the Cancer Center Support Grant P30CA016087 at the Laura and Isaac Perlmutter Cancer Center. This work was supported by the Irvington Institute fellowship program of the Cancer Research Institute (M. X.); Training program in Immunology and Inflammation 5T32AI100853 (M.P.); the Helen and Martin Kimmel Center for Biology and Medicine (D.R.L.); the Colton Center for Autoimmunity (D.R.L.); and National Institutes of Health grant R01DK103358 (R.B. and D.R.L.). D.R.L. is an Investigator of the Howard Hughes Medical Institute.

## Author Contributions

M.X. and M.P. designed and performed all experiments and analyzed the data. Y.D. performed blinded histology scoring on colitis sections. C.A and C.G. assisted with *in vivo* and *in vitro* experiments. R.Y. and M.P. performed RNA-seq analysis. R.B. supervised RNA-seq analysis. M.X., M.P., and D.R.L. wrote the manuscript with input from the co-authors. D.R.L. supervised the research and contributed to experimental design.

## EXTENDED DATA FIGURE LEGENDS

**Extended Data Figure 1:**
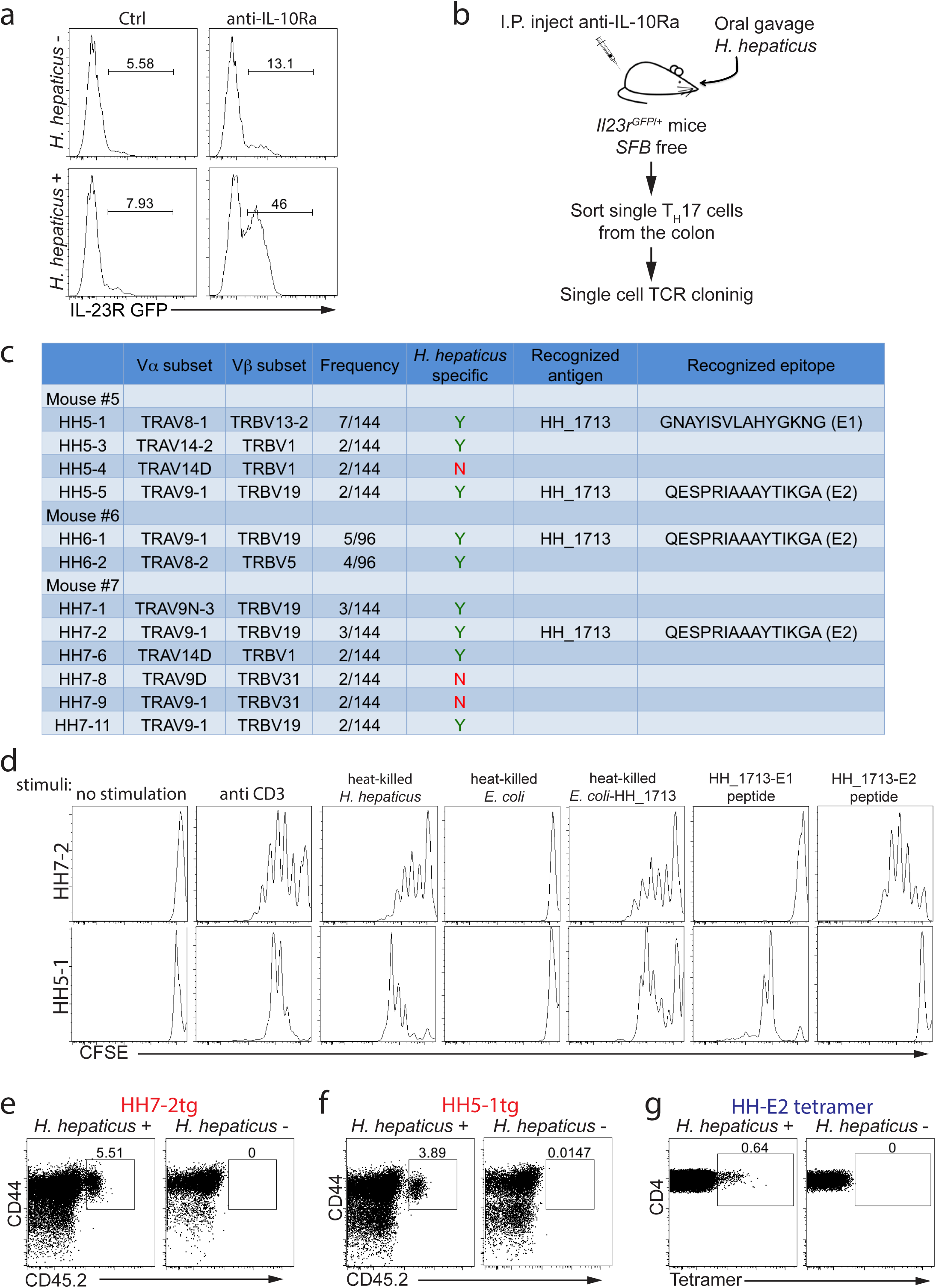
Cloning and characterization of *H. hepaticus*- specific T_H_17 TCRs, and generation of TCR transgenic (TCRtg) mice and MHC-II tetramers. **a,** IL-23R-GFP expression in CD4^+^ T cells from the large intestines of mice with and without *H. hepaticus* colonization and after IL-10Ra blockade. **b,** Experimental scheme for cloning *H. hepaticus*-induced single T_H_17 cell TCRs under IL-10Ra blockade. **c,** Summary of the twelve dominant *H. hepaticus*-induced T_H_17 TCRs. **d,** *In vitro* activation of CFSE-labeled naive HH7-2tg and HH5-1tg cells by indicated stimuli in the presence of antigen-presenting cells. **e,** Expansion of donor-derived HH7-2tg (CD45.2) cells in the large intestine (LI) of *H. hepaticus*-colonized or -free CD45.1 mice, gated on total CD4^+^ T cells. **f,** Expansion of donor-derived HH5-1tg (CD45.2) cells in the LI of *H. hepaticus*-colonized or -free (CD45.1) mice, gated on total CD4^+^ T cells. **g,** HH-E2 tetramer staining of CD4^+^ T cells from the LI of *H. hepaticus*-colonized or -free mice.

**Extended Data Figure 2:**
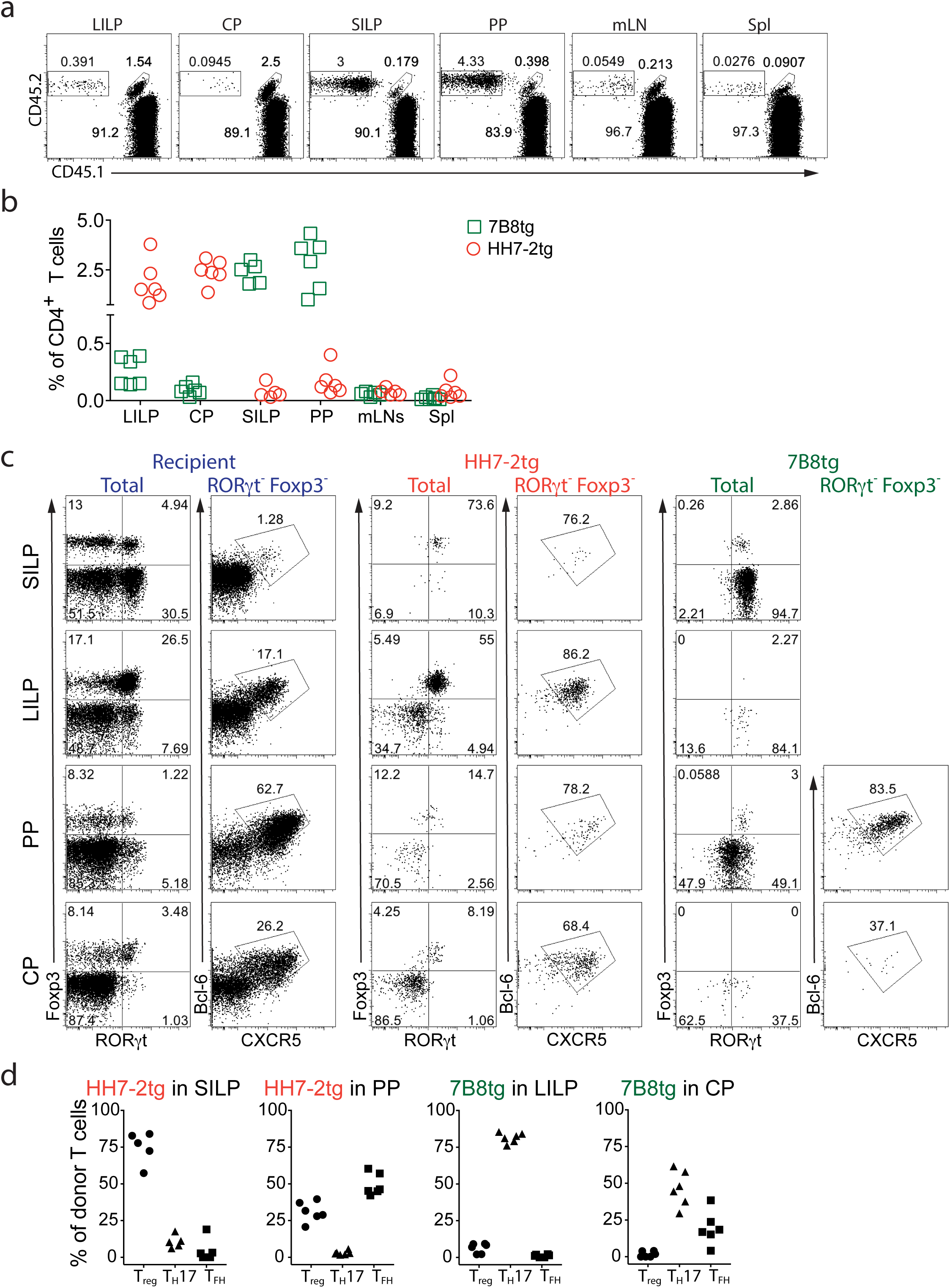
Enrichment and differentiation of HH7-2tg and 7B8tg T cells in distinct anatomical sites in WT recipient mice colonized with SFB and *H. hepaticus*. **a,** Representative flow cytometry plots of donor-derived HH7-2tg (CD45.1/45.2) and 7B8tg (CD45.1/45.1) T cells in indicated tissues of mice colonized with SFB and *Helicobacter hepaticus*, gated on total CD4^+^ T cells. **b,** Proportions of donor-derived HH7-2tg and 7B8tg T cells among total CD4^+^ T cells in indicated tissues. Data are from one of 3 experiments, with total of 15 mice in the 3 experiments. **c,** Representative flow cytometry plots of RORγt, Foxp3, Bcl-6 and CXCR5 expression in CD4^+^ T cells from the host and from HH7-2tg and 7B8tg donors in different tissues (n=15). **d,** Frequencies of T_reg_ (Foxp3^+^), T_H_17 (Foxp3^-^RORγt^+^) and T_FH_ (Bcl-6^+^CXCR5^+^) cells in donor-derived HH7-2tg and 7B8tg cells in different tissues. Data are from one of 3 experiments, with total of 15 mice in the 3 experiments. SILP: small intestinal lamina propria; LILP: large intestinal lamina propria; PP: Peyer's patches; CP: cecal patches; mLNs: mesenteric lymph nodes; and Spl: spleen.

**Extended Data Figure 3:**
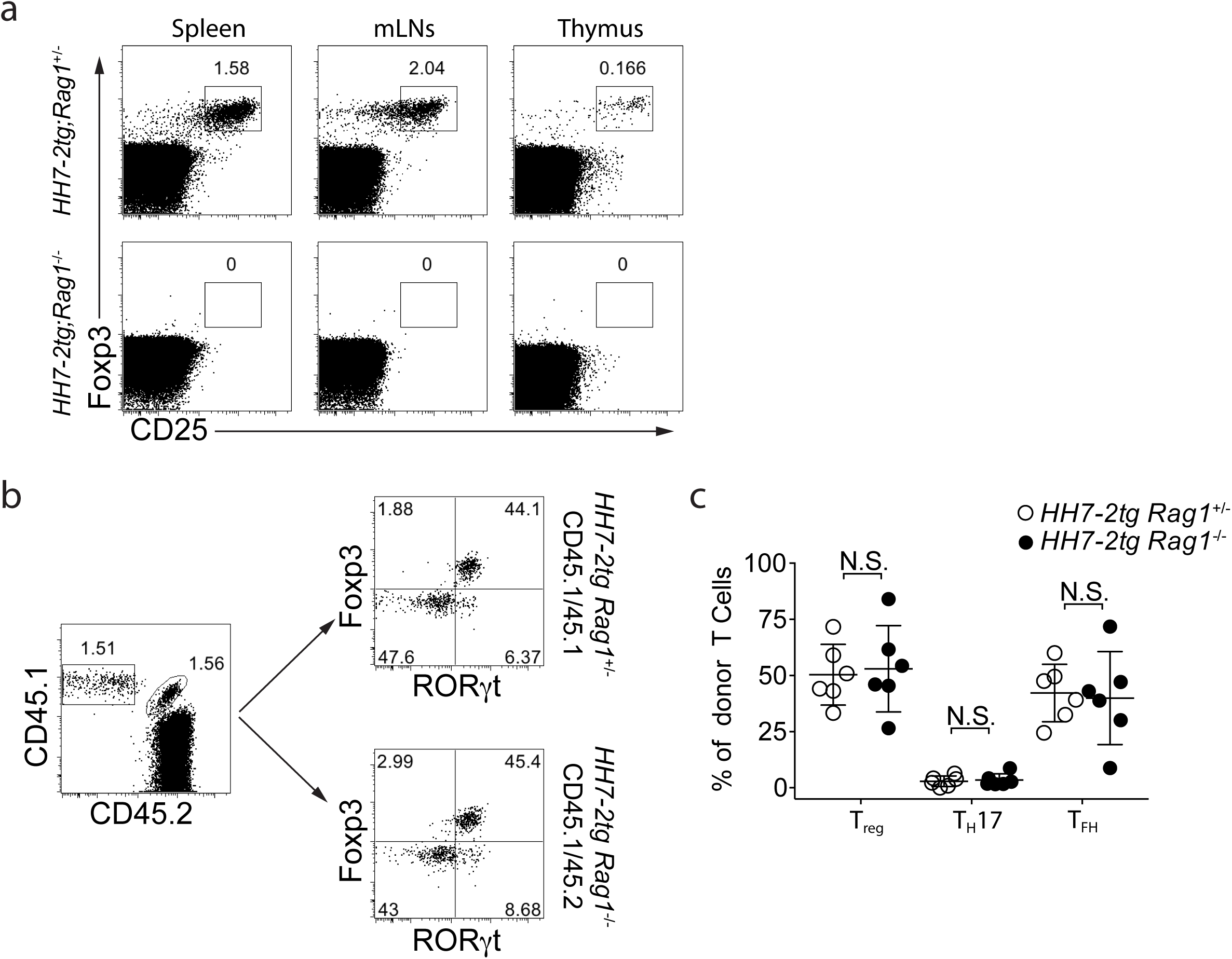
Characterization of HH7-2tg;*Rag*1^-/-^ mice. **a,** Representative flow cytometry plots of T_reg_ (Foxp3^+^CD25^+^) frequency in indicated tissues of *H. hepaticus*-free HH7-2tg;*Rag1*^+/-^ (n=3) or HH7-2tg;*Rag1*^-/-^ (n=3) mice. **b,** Development of co-transferred congenic isotype-labeled HH7-2tg *Rag1*^+/-^ (CD45.1/45.1) and *Rag1*^-/-^ (CD45.1/45.2) naïve T cells in the LI of *H. hepaticus*-colonized WT B6 mice. Representative flow cytometry plots of donor and recipient T cell frequency (left), and RORγt and Foxp3 expression (right). **c,** Frequencies of T_reg_ (Foxp3^+^), T_H_17 (Foxp3^-^RORγt^+^) and T_FH_ (Bcl-6^+^CXCR5^+^) cells among HH7-2tg *Rag1*^+/-^ (CD45.1/45.1) (n=6) and *Rag1*^-/-^ (CD45.1/45.2) (n=6) donor-derived T cells. Error bars: mean ± 1 SD. Statistics were calculated by unpaired *Welch t-*test, N.S. (P≥0.05, not significant).

**Extended Data Figure 4:**
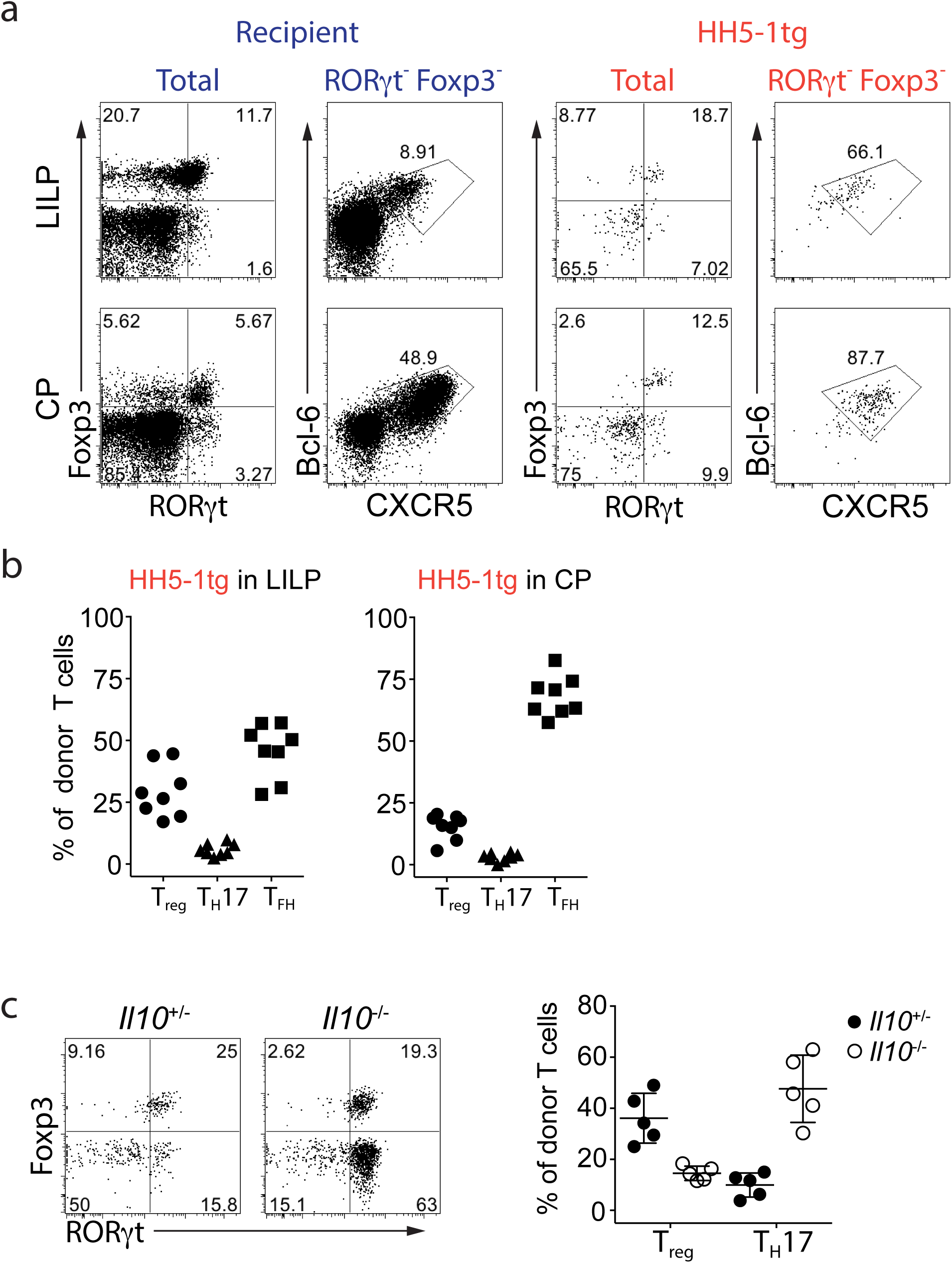
Differentiation of adoptively transferred naïve HH5-1tg T cells in *H. hepaticus*-colonized mice. **a,b,** 2000 naïve HH5-1tg cells (CD45.1/45.2) were adoptively transferred into WT B6 mice (CD45.2/45.2) colonized with *H. hepaticus*. Cells from LI and CP were analyzed two weeks after transfer. Representative flow cytometry plots are shown for RORγt, Foxp3, Bcl-6 and CXCR5 expression in donor-derived and recipient CD4^+^ T cells in indicated tissues. Frequencies of T_reg_ (Foxp3^+^), T_H_17 (Foxp3^-^RORγt^+^) and T_FH_ (Bcl-6^+^CXCR5^+^) among HH5-1tg donor T cells (n=8). Data are a summary of two independent experiments. **c,** Naïve HH5-1tg cells were adoptively transferred as above into *Il10*^+/-^ and *Il10*^-/-^ mice (CD45.2/45.2) colonized with *H. hepaticus*. Cells from LI were analyzed two weeks after transfer (n=5). Representative flow cytometry plots of RORγt and Foxp3 expression in HH5-1tg donor cells in LILP (left), and a compilation of frequencies of T_reg_ (Foxp3^+^) and T_H_17 (Foxp3^-^RORγt^+^) among HH5-1tg donor cells (right). Error bars: mean ± 1 SD. Statistics were calculated by unpaired *Welch t-*test, ** p<0.01, *** p<0.001.

**Extended Data Figure 5:**
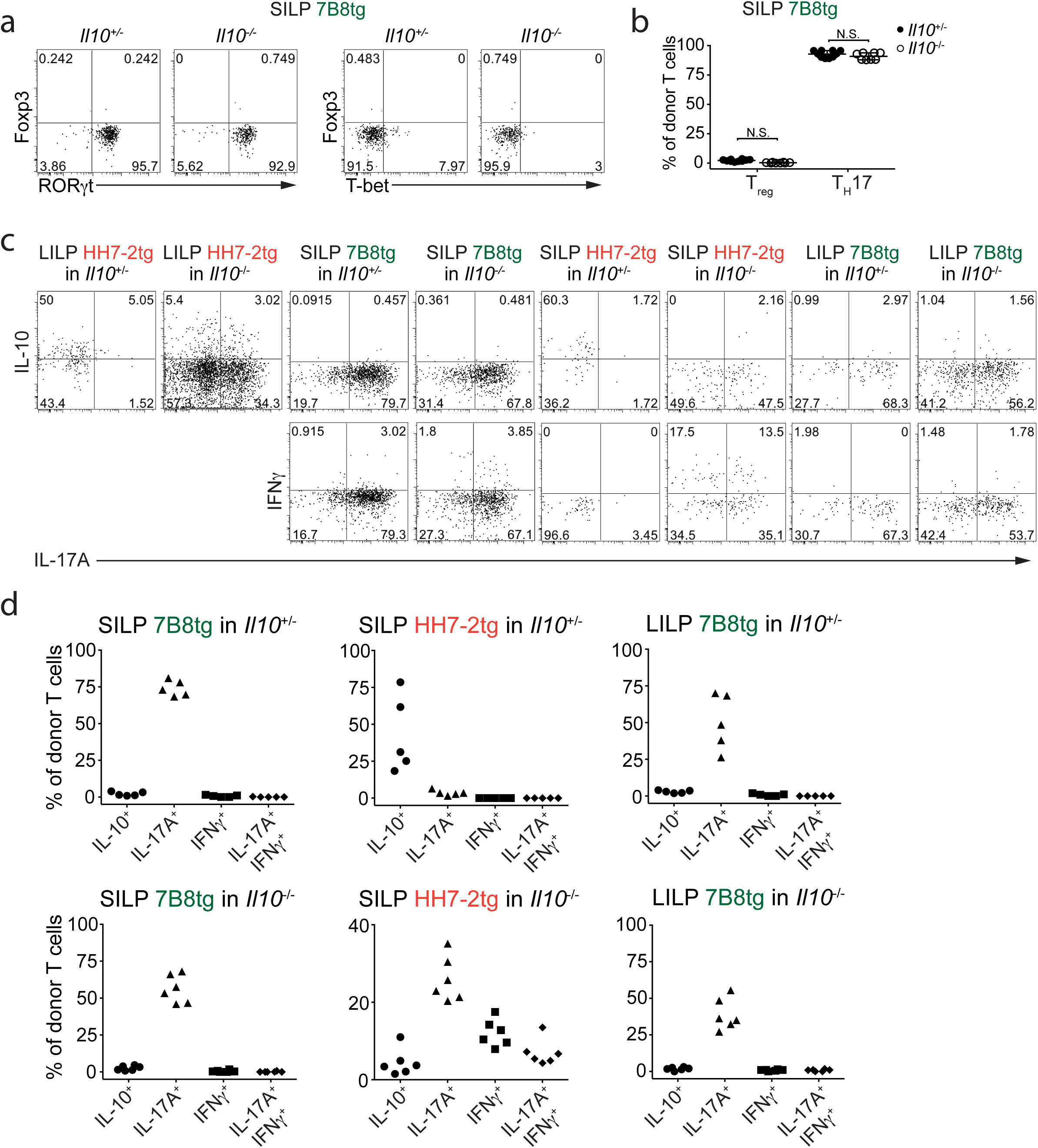
Differentiation of transferred HH7-2tg and 7B8tg T cells in *Il10*^+/-^ and *Il10*^-/-^ mice. **a,** Representative flow cytometry plots of Foxp3, RORγt and T-bet expression in 7B8tg cells in the SILP of *Il10*^+/-^ (n=10) and *Il10*^-/-^ (n=8) recipient mice. **b,** Frequencies of T_reg_ (Foxp3^+^) and T_H_17 (Foxp3^-^RORγt^+^) cells among SILP 7B8tg donor-derived cells in *Il10*^+/-^ (n=10) and *Il10*^-/-^ (n=8) mice. Data are a summary of four independent experiments. Error bars: mean ± 1 SD. Statistics were calculated by unpaired *Welch t-*test, N.S. (P≥0.05, not significant). **c,** Representative flow cytometry plots of IL-10, IL-17A and IFNγ expression in transferred 7B8tg and HH7-2tg cells from LILP and SILP of *Il10*^+/-^ and *Il10*^-/-^ mice after re-stimulation (n=5 or 6). **d,** Proportions of transferred 7B8tg and HH7-2tg cells in the SILP and LILP of *Il10*^+/-^ and *Il10*^-/-^ mice that express IL-10, IL-17A and IFNγ after re-stimulation (n=5 or 6). Data are a summary of two independent experiments.

**Extended Data Figure 6:**
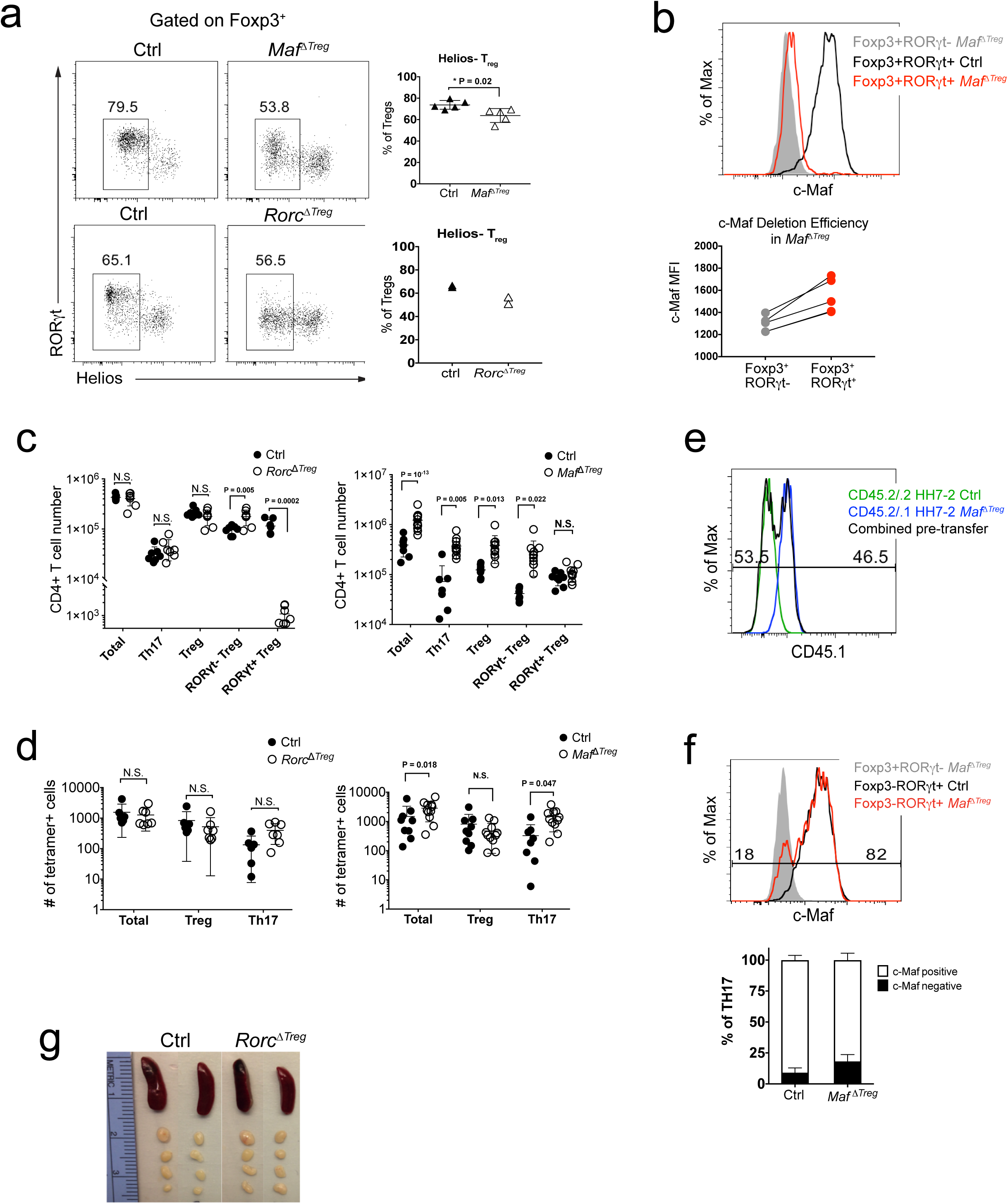
Extended characterization of *Maf* ^ΔTreg^ and *Rorc*^ΔTreg^ animals. **a,** Left, representative flow cytometry plots of RORγt and Helios expression in the T_reg_ compartment. Right, summary of frequencies from two independent experiments. Error bars: mean ± 1 SD. Statistics were calculated by unpaired *Welch t-*test. **b,** Incomplete depletion of c-Maf protein in RORγt^+^ Tregs in *Maf* ^ΔTreg^ mice shown by a representative flow cytometry graph (upper) and a compilation of frequencies (lower). **c,** Absolute numbers of indicated CD4^+^ T cell populations in the LILP of *Rorc*^ΔTreg^ (left) and *Maf* ^ΔTreg^ (right) mice. Data are a summary of 3 independent experiments for *Rorc*^ΔTreg^ (n=7) and littermate controls (n=7) and 4 independent experiments for *Maf* ^ΔTreg^ (n=11) and littermate controls (n=8). Error bars: mean ± 1 SD. Statistics were calculated by unpaired *Welch t-* test, N.S. (P≥0.05, not significant), * p<0.05, ** p<0.01, *** p<0.001. **d,** Absolute numbers of indicated HH-E2 tetramer+ T cell populations in the LILP of *Rorc*^ΔTreg^ (left) and *Maf* ^ΔTreg^ (right) mice. Data are a summary of 3 independent experiments for *Rorc*^ΔTreg^ (n=7) and littermate controls (n=6) and 4 independent experiments for *Maf* ^ΔTreg^ (n=11) and littermate controls (n=8). Error bars: mean ± 1 SD. Statistics were calculated by unpaired *Welch t-*test, N.S. (P≥0.05, not significant), * p<0.05, ** p<0.01, *** p<0.001. **e,** Flow cytometry plot depicting ratio of co-transferred cells. **f,** Representative flow cytometry plot of c-Maf expression in T_H_17 cells (Foxp3^-^ RORγt^+^) from LILP of control (black) and *Maf* ^ΔTreg^ (red) mice (left). The c-Maf negative population is defined by gating on Foxp3^+^RORγt^-^ T_reg_ from *Maf* ^ΔTreg^ mice (solid grey). Below, summary of frequencies of c-Maf expression in Th17 cells for control (n=6) and for *Maf* ^ΔTreg^ (n=9) mice from 3 independent experiments. **g,** Spleens and mLNs of *Rorc*^ΔTreg^ and control mice colonized by *H. hepaticus* for 5-6 weeks.

**Extended Data Figure 7:**
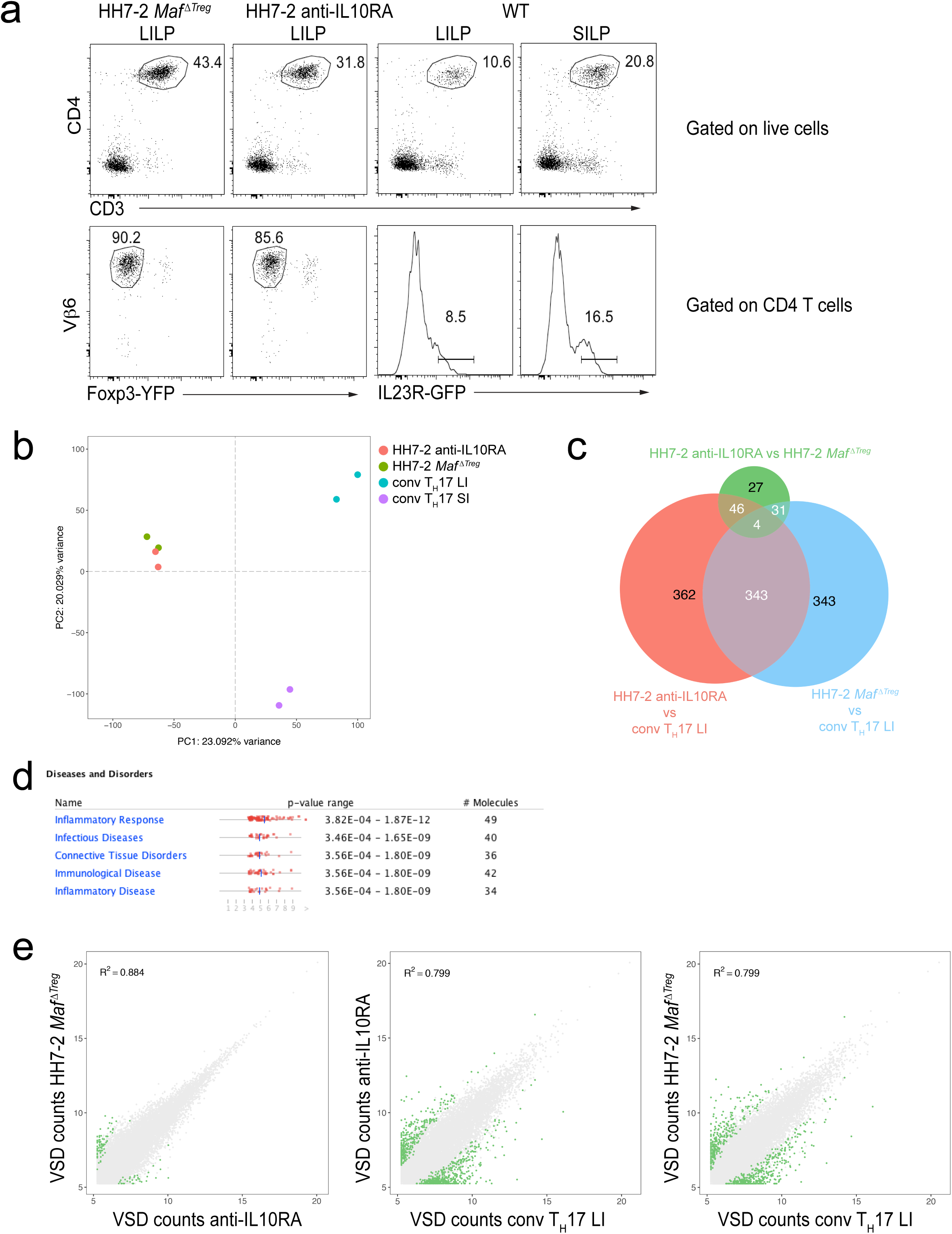
Transcriptional profiling of conventional and *H. hepaticus*-specific T effector cells. **a,** Flow cytometry analysis of donor-derived HH7-2tg T effector cells from *H. hepaticus*-colonized mice and conventional IL-23R-GFP^+^ T_H_17 cells from SFB-colonized mice. Gates in the lower panel were used for sorting to perform RNA-seq. **b,** Principal component analysis of RNA-seq data from sorted cell populations. **c,** Venn diagram depicting differentially expressed genes (FDR < 0.1) between indicated comparisons identified by DESeq2. **d,** Significantly enriched disease pathways in the set of 149 shared genes upregulated in HH7-2tg *Maf* ^ΔTreg^ and HH7-2tg anti-IL-10ra compared to conventional LI T_H_17. **e,** Scatter plots of Variance Stabilized Data (VSD) counts. Green dots represent differentially expressed genes identified by DESeq2 (FDR < 0.1). R^2^ calculated by linear regression.

**Extended Data Figure 8:**
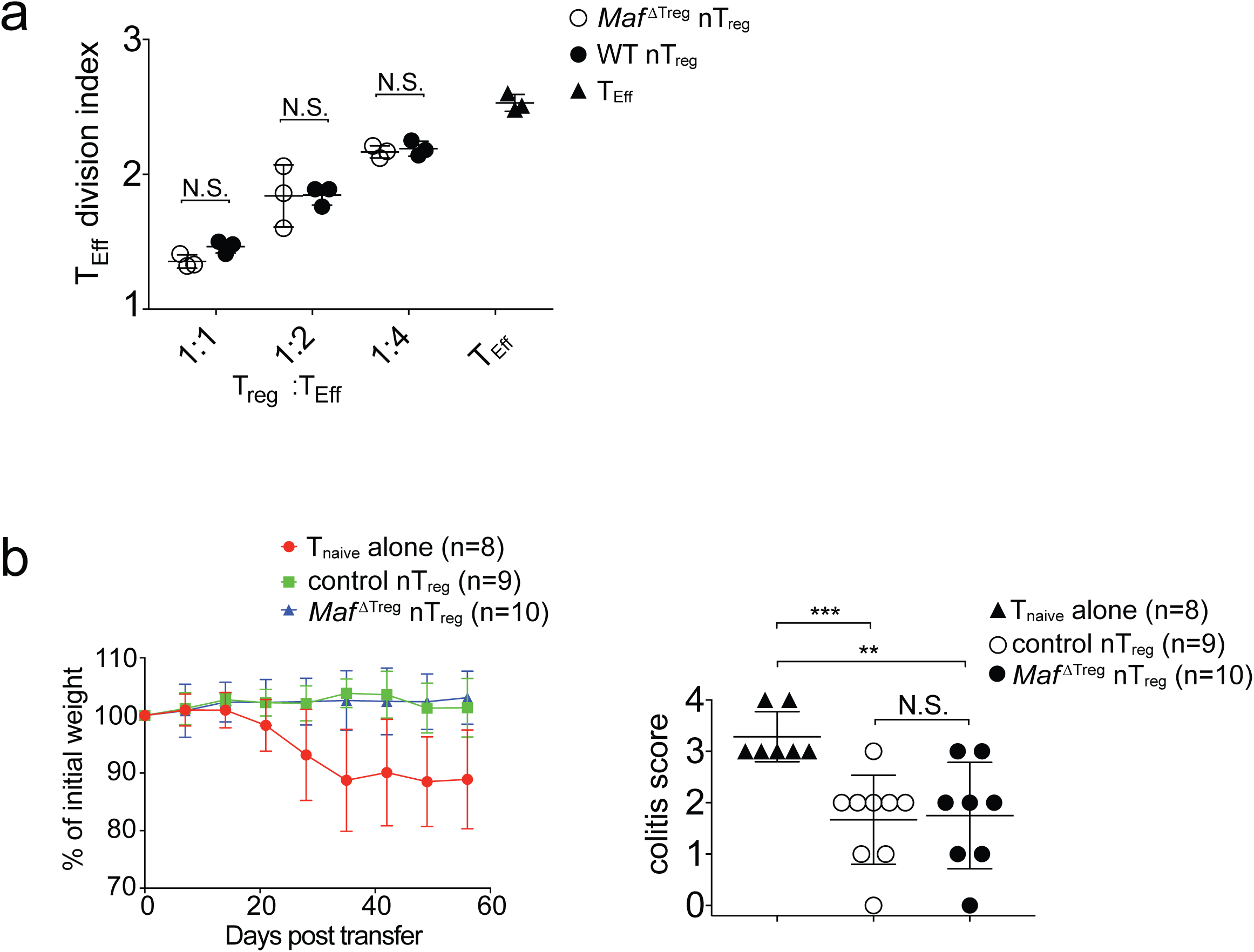
c-Maf-deficient nT_reg_ cells retain suppressive function. **a,** Equivalent inhibitory function of nT_reg_ cells from *Maf* ^ΔTreg^ and control mice in the *in vitro* proliferative response of CD4^+^ T cells (T_Eff_). Three data points are from one of two independent replicates. **b,** Activity of nTreg cells in the transfer-mediated colitis model. Percentage weight change and colitis histology scores (right) of *Rag*1^-/-^ mice adoptively transferred with naïve T cells alone (n=8), or naïve T cells in combination with nT_reg_ cells from *Maf* ^ΔTreg^ (n=10) or control (n=9) mice. Data are a summary of two independent experiments. Statistics were calculated by unpaired *Welch t-* test, N.S. (P≥0.05, not significant), ** p<0.01, *** p<0.001.

